# The neuronal ceroid lipofuscinosis protein Cln7 regulates neural development from the post-synaptic cell

**DOI:** 10.1101/278895

**Authors:** Kyle J. Connolly, Megan B. O’Hare, Alamin Mohammed, Katelyn M. Aitchison, Niki C. Anthoney, Matthew J. Taylor, Bryan A. Stewart, Richard I. Tuxworth, Guy Tear

## Abstract

The neuronal ceroid lipofuscinoses (NCLs) are a group of fatal, monogenic neurodegenerative disorders with an early onset in infancy or childhood. Despite identification of the genes disrupted in each form of the disease, their normal cellular role and how their deficits lead to disease pathology is not fully understood. Cln7, a major facilitator superfamily domain-containing protein, is affected in a late infantile-onset form of NCL. Using the *Drosophila* larval neuromuscular junction as a model to study neural development, we demonstrate that Cln7 is required for the normal growth of synapses. In a *Cln7* mutant, synapses fail to develop fully leading to reduced function and behavioral changes with dysregulation of TOR activity. *Cln7* expression is restricted to the post-synaptic cell and the protein localizes to vesicles immediately adjacent to the post-synaptic membrane. Our data suggest an involvement for Cln7 in regulating trans-synaptic communication.

## Introduction

The study of early-onset, inherited forms of neurodegenerative disease provides an opportunity to identify how single gene defects lead to neurodegeneration. The neuronal ceroid lipofuscinoses (NCLs) are a group of such monogenetic neurodegenerative disorders with disease onset early in infancy or childhood (5-8 years in the most common form but congenital and rare adult-onset forms also exist). Patients present with a loss of visual acuity followed by onset of seizures and psychiatric disturbances. A progressive mental decline follows, accompanied by loss of motor skills and death usually occurs by the age of 20-30 (1). The NCLs share a hallmark pathology: accumulation of fluorescent storage material indicating lysosomal dysfunction. The progression of NCL pathology appears similar to many other neurodegenerative disorders in that changes to synapses occur very early and a local activation of glia precedes a regionally specific loss of neurons (2, 3). Their early onset may well indicate that one or more key cellular pathways are impaired directly, rather than a consequence of the progressive buildup of factors as may occur in the late-onset diseases. There is also likely to be a significant developmental component to the neuropathology, potentially pre-disposing the nervous system to degeneration.

The NCLs are recessively inherited monogenic disorders (with the exception of one rare adult-onset autosomal dominant form) and mutations have been identified in 14 genes responsible for NCL with varying ages of onset. A limited set of these genes is conserved in *Drosophila* suggesting that they are performing core functions necessary for normal neuronal health. Mutations in *CLN7/MFS-domain containing 8* (*CLN7*/*MFSD8*) are responsible for late-infantile onset NCL (LINCL or CLN7 disease), with disease onset at 1.5-5 years of age (4). The CLN7 protein is predicted to be a member of the major facilitator superfamily of transporters, the majority of which have twelve membrane spanning domains (5). In cell culture experiments with tagged forms, CLN7 protein is localized primarily in lysosomes (5-7) and has been identified by lysosomal proteomics (8). Mouse embryonic fibroblasts derived from *CLN7* deficient mice show a loss of lysosomal proteins and deficits in mTOR reactivation (9). Consistent with this, phenotypes in mutant mice show lysosomal dysfunction and a potential impairment of autophagy (10). However, its *in vivo* function remains unknown.

*CLN7* is expressed widely within the murine CNS with higher mRNA expression in the cerebellum and hippocampus (7). A Cln7-lacZ reporter construct also reveals expression in the entire cerebral cortex, caudate putamen and thalamus. In the spinal cord *CLN7* is expressed in the gray matter but excluded from white matter (11). Recent single cell RNA transcriptomics from human brain cells indicate that *CLN7* is expressed in neurons and oligodendrocytes lineages but is largely undetectable in astrocytes (12), http://scope.aertslab.org/. We similarly find that *Drosophila Cln7* is expressed in the CNS. However, using YFP-tagged Cln7 expressed from its endogenous locus, we observe that neural expression of *Cln7* is largely restricted to the optic system but that *Cln7* is more widely expressed in glia (13), including in the glia that form the blood-brain-barrier and in the ensheathing glia that wrap axons (Fig. 1). *Cln7* is also expressed in the neuromuscular system where high levels are found in the postsynaptic muscles. This corresponds with co-localisation of Cln7 protein with the post-synaptic density protein, PSD-95, in the outer plexiform layer of the murine retina (14).

**Figure 1.**
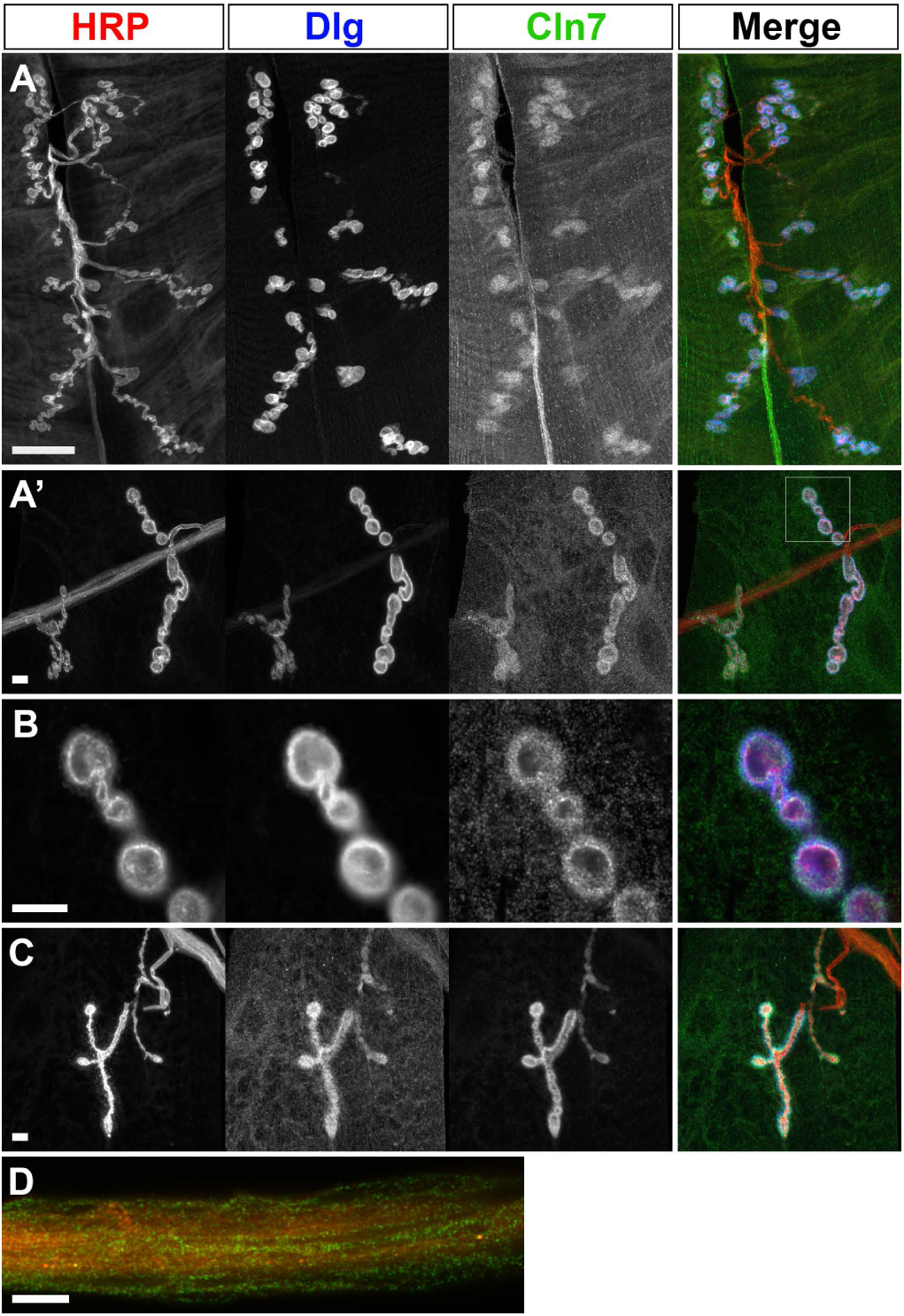
Cln7 localises to a vesicular compartment at the post-synaptic membrane. A and B). Late larval NMJs from a YFP-Cln7 knock-in reporter line fixed and stained for the neuronal plasma membrane (anti-HRP, red), the sub-synaptic reticulum membrane (anti-Dlg, blue) and for YFP-Cln7 (anti-GFP, green). YFP-Cln7 is restricted to the post-synaptic muscle cell at the larval neuromuscular junction. A shows NMJ6-7 and B shows NMJ4b (B). The boxed region in B was imaged at higher resolution in B’. Distinct Cln7^+^ vesicles are concentrated within and around the sub-synaptic reticulum. C). Cln7 does not require the endogenous 3’ UTR for recruitment to the synapse. A Myc-BioID-Cln7 fusion protein is still recruited to the sub-synaptic reticulum when overexpressed from a cDNA construct in the muscle lacking the endogenous 3’UTR. Anti-Myc (green) was used to visualize the Cln7 fusion. D). YFP-Cln7 is also expressed in the enseathing glia that wrap axons in the nerves. Transverse optical section through intersegmental nerve 1b stained with anti-HRP (neurons, red) anti-GFP (YFP-Cln7, green). Bar = 10 μm (A and D); 5 μm (B and C).

Evidence is building in several models that loss of the *CLN* genes impacts on synapse function, suggesting that they play a common role either within neurons or non-neuronal cells in the development or maintenance of synapses (2, 15, 16). Since neurodevelopmental defects (unless severe) are difficult to identify in standard rodent models, we turned to the *Drosophila* neuromuscular junction (NMJ), which provides a simple, genetically tractable model system to study the development and maturation of neurons and the assembly and function of synapses. Many genes associated with human neurological conditions, including developmental disorders and neurodegenerative diseases, have roles regulating NMJ development (7, 17). As the *Drosophila* larva proceeds through development, increasing 100-fold in volume over 3-days, the concomitant expansion of the NMJ is regulated by homeostatic processes to maintain the correct levels of innervation to the body wall muscles. Multiple cellular pathways mediate both short-term and longer-term homeostasis, acting at both sides of the synapse, and many of these mechanisms also act similarly to regulate homeostasis in mammalian neurons (18). In addition, normal autophagic flux and endo-lysosomal function are required to maintain the synapse with mutants affecting lysosomal function, including Spinster and the cation channel Trpml, and autophagy, including several of the *atg* mutants, impacting on NMJ development and expansion of the pre-synaptic compartment (19-21).

To study a potential role for Cln7 *in vivo*, we generated a *Drosophila* model for CLN7 disease by deleting the N-terminal half of the *Cln7* gene. While the animals remain viable, they display impaired locomotion and an associated reduction in synapse size and altered synapse function. We show that a population of Cln7-containing vesicles localizes at the post-synaptic side of the synapse but autophagy is largely intact in the *cln7* mutants and retrograde BMP signaling pathway driving pre-synapse growth functions normally. We show that Cln7 can be purified in a complex with the TOR complex activator, Rheb, which suggests Cln7 regulates neural development via TOR from a post-synaptic vesicular compartment.

## Results

### Generation of a *Drosophila* model of CLN7 disease

Disruption of the *MFSD8/CLN7* gene has been identified as the cause of the LINCL form of the NCLs (4). Subsequently, more than 30 different *Cln7* mutations have been characterized. The *Drosophila* gene CG8596 shows 57% amino acid homology to human *CLN7* and is likely to be the unique *Drosophila* orthologue of *Cln7*. The gene encodes a predicted 12-spanning transmembrane protein that has strong sequence homology to the multifacilitator family of solute transporters (22) (Supp. Fig. 1A). To study the functional role of Cln7, we generated two loss-of-function alleles of *Cln7* by imprecise excision of the P-element NP0345 inserted in the upstream region proximal to the start codon (Supp. Fig. 1B). Two alleles were generated. Exons 1-5 were excised with some Pelement sequence remaining in the *Cln7*^*84D*^ allele, and exons 1 and 2 and part of exon 3 were excised in the *Cln7*^*36H*^ allele (Supp. Fig. 1B). Both mutant alleles are viable and fertile when homozygous.

### Cln7 is localized to the post-synaptic side of the neuromuscular junction

In a parallel study we generated Venus-YFP knock-in forms of Cln7 by CRISPR/Cas9-mediated gene editing to act as a reporter of expression and sub-cellular localization (13). We identified that Cln7 is expressed in neurons in the visual system but elsewhere in the CNS is primarily a glial protein in the CNS. It is also expressed in the body wall muscles that form the post-synaptic cells of the neuromuscular junction (NMJ) (13). We further examined the localization of Cln7 using higher-resolution Airyscan confocal imaging. We find that Cln7 is present isolated vesicles throughout the muscle cell but also concentrates at the post-synaptic site (Fig. 1A). Surrounding each pre-synaptic swelling (or bouton) is a convoluted membranous structure known as the sub-synaptic reticulum (SSR). The SSR contains the post-synaptic glutamate receptors. The commonly-used marker of the SSR, anti-Discs large (Dlg, the PSD-95 ortholog), appears in a continuous pattern around the boutons but higher-resolution imaging reveals that Cln7 is clearly localized to discrete vesicles immediately surrounding the post-synaptic site, rather than in the plasma membrane of the cell (Fig. 1B). Recruitment of Cln7 to the synapse is maintained even when a Cln7 fusion protein (Myc-BioID-Cln7) is overexpressed under the control of Mef2-Gal4 (Fig. 1C). The nature of the Cln7^+^ vesicles is not clear but they do not apper to contain Cathepsin L, a lysosomal marker, and lysosomes do not concentrate at the synapse (not shown).

### Cln7 is required for normal synaptic growth

The distribution of Cln7 within post-synaptic vesicles suggests the protein may function to regulate the synapse and, since LINCL may be a result of synapse failure, we investigated the consequence of a loss of *Cln7* function.

The *Drosophila* NMJ is a readily accessible glutamatergic synapse and an excellent model system to study synapse function, so this discovery afforded us an opportunity to ask how Cln7 might function at the post-synaptic site. The peripheral NMJs in late stage larvae are easily identifiable using immunofluorescence and the innervations are simple, with 2-4 individual motoneurons contacting each muscle cell. The type I motoneurons that innervate the muscle can be further subdivided into two classes called type Ib and type Is classified according to the size of the presynaptic terminal swellings known as boutons (Fig. 2A). The type Ib terminals provide tonic stimulation and the type Is phasic (23). The overall size of the NMJ is governed by intra-neuronal and trans-synaptic homeostatic mechanisms that modulate NMJ size during development to match pre-synapse size and strength to muscle size (18). Many mutations affecting the structural and electrical properties of the NMJ consequently affect its size at the late larval stage, including many mutations related to human neurological diseases (17).

**Figure 2.**
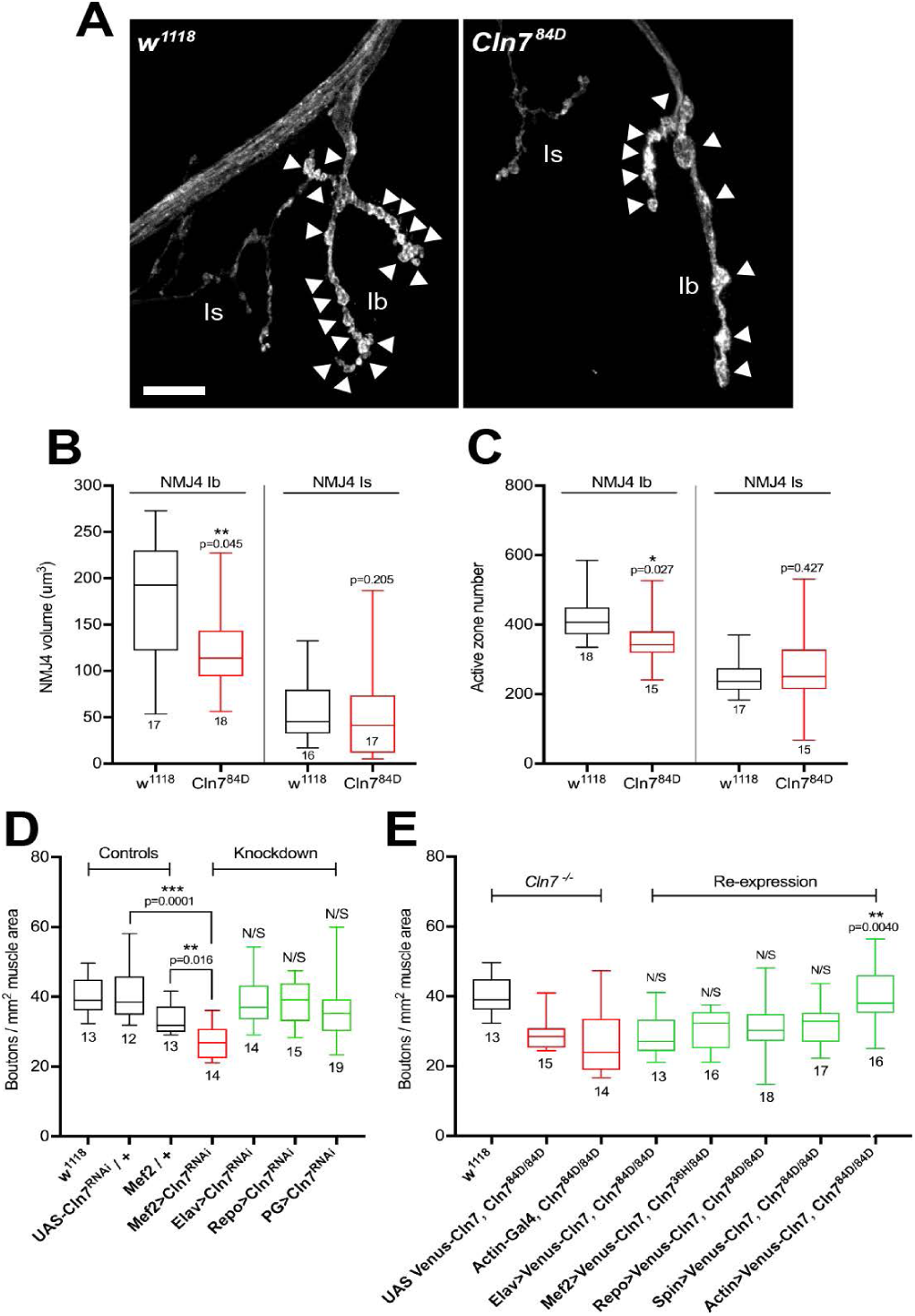
Loss of *Cln7* leads to neurodevelopmental defects. A). Anti-HRP staining of the late larval NMJ of *w*^*1118*^ control and *Cln7*^*84D*^ flies to visualize the neuronal membrane. Swellings of the pre-synaptic membrane (boutons) are indicated with arrowheads. There are fewer boutons in *Cln7*^*84D*^ larvae but these are occasionally swollen. Scale bar = 10 μm. B). The volume of type Ib junction is reduced in *Cln7*^*84D*^ flies but the type Is junction is unchanged. C). Active zones number is reduced in *Cln7*^*84D*^ type Ib but not in type Is junctions. (B&C; 2-tailed t-tests, n is indicated). D). The number of boutons can be used as a proxy for junction size. Type Ib junctions are smaller when *Cln7* expression is knocked down post-synaptically in muscle (Mef2-gal4) but not with pre-synaptic (Elav), pan-glial (Repo) or perineurial glial knockdown (PG). E). Normal NMJ development is restored only when Cln7 is re-expressed ubiquitously (actin-Gal4) in the mutant background. (D&E: ANOVA with Dunnett’s post-hoc comparisons, n is indicated).

We find that in *Cln7*^-/-^ animals the NMJ is reduced in size. Manual counting of the swellings of the terminal (boutons) is commonly used as a simple measure of terminal size as the boutons contain most of the active zone synaptic vesicle release sites. At muscle 4 we see a significant reduction of the type Ib motoneuron terminals in the *Cln7* mutant larvae when compared to isogenic control larvae (Fig. 2A). Some of the boutons in the *Cln7* larvae appeared larger so we also used a non-subjective method of measuring terminal volume from 3D-rendered confocal images. Again, the type Ib terminals were significantly smaller than controls but, interestingly, the type Is terminals were unaffected (Fig. 2B). As a further measure of synapse size we counted the number of active zones at the NMJ. Active zones are visualized with an antibody to Bruchpilot (ELKS), an active zone component (24). Bruchpilot positive puncta were counted at muscle 4 and this confirmed that the number of active zones within Ib boutons is reduced in *Cln7*^-/-^ animals concomitant with the reduction in volume but is unchanged in type Is junctions (Fig. 2C). Both alleles of CLN7 had an identical phenotype as does the transheterozygous combination. Interestingly, both alleles were also haploinsufficient, suggesting some processes important for NMJ development are critically dependent on the levels of the CLN7 protein: potentially the stoichiometry of a membrane complex. These results suggest a requirement for Cln7 for normal development of the NMJ.

### Cln7 function is required in the post-synaptic side of the NMJ

To investigate whether Cln7 is required in the post synaptic cell we used RNAi to reduce *Cln7* expression in various cell types at the synapse. Standard GAL4 drivers were used to express dsRNA specifically targeted at *Cln7* either pre-synaptically (Elav-Gal4), post-synaptically in the muscle (Mef2-Gal4), in all glia (Repo-Gal4) and in only the perineurial glia (PG-Gal4) that form the blood-brain-barrier, where *Cln7* is also highly expressed (13). Only knockdown of *Cln7* in the muscle replicated the under-development phenotype at the NMJ (Fig. 2D), consistent with the expression of *Cln7*. We also performed the converse experiment of re-expressing YFP-Cln7 in the *Cln7*^-/-^ background, again with ubiquitous expression (actin-Gal4) or expression restricted to neurons (Elav-Gal4), to muscle (Mef2-Gal4), in both neurons and muscle together (Spin-Gal4) or to glia (Repo-Gal4). Muscle expression with Mef2- or Spin-Gal4 was only capable of partial rescue of function which narrowly failed to reach significance. Only ubiquitous expression fully rescued the phenotype (Fig. 2E) indicating that some Cln7 function is required elsewhere in addition to the muscle.

### Loss of Cln7 leads to synapse dysfunction

To examine the consequence of the abnormal synaptic growth we characterized synaptic physiology and larval motility. An electrophysiological analysis of the NMJ was carried out using intracellular recordings. We recorded the spontaneous transmitter release at the larval NMJs for two minutes to calculate miniature excitatory junction potential (mEJP) frequency and amplitude in 0.5, 0.75, 1.0 and 2.0 mM extracellular Ca^2+^ concentration but there was no significant difference due to genotype in either case (Fig. 3A; 2-way ANOVA with Sidak’s multiple comparison test, p=0.483). Taken together, we concluded there was likely to be little overt change in spontaneous synaptic vesicle release or post-synaptic receptor density. Consistent with this, there were no overt changes in the levels of the Glutamate receptors, GluRIIA or GluRIII, in *Cln7* mutant boutons when compared to controls (not shown).

**Figure 3.**
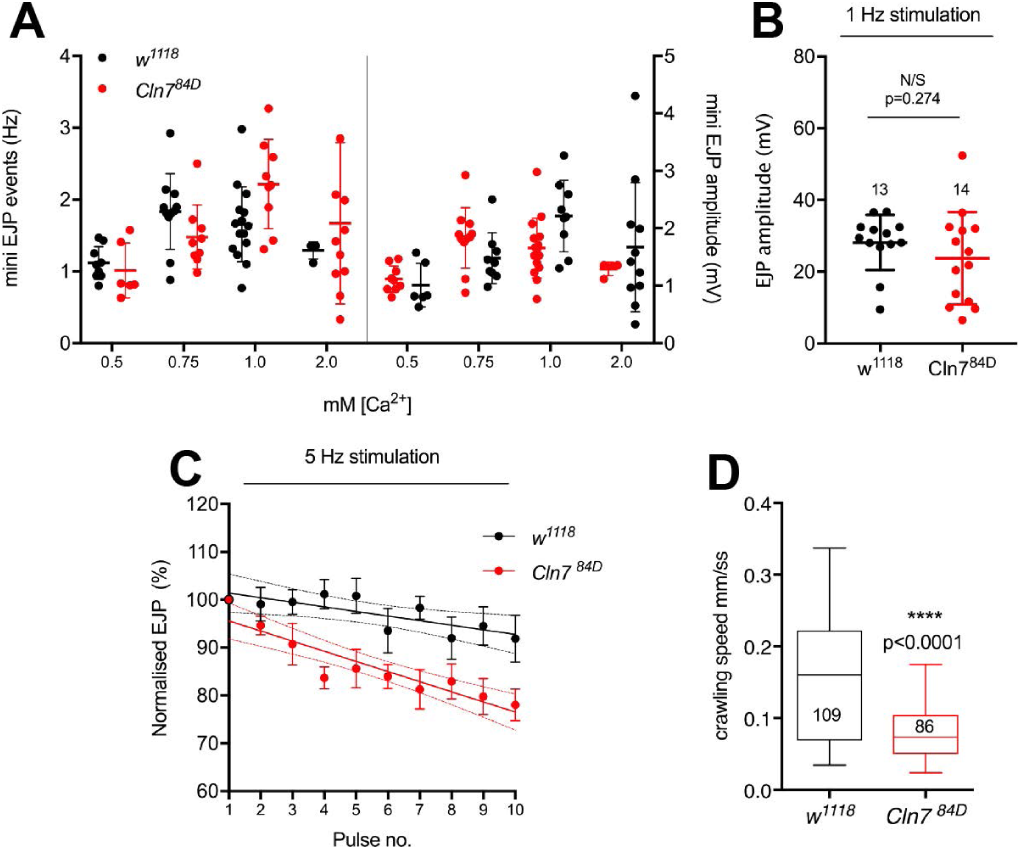
Altered neural function in *Cln7* mutant flies. A-C). Electrophysiological recordings from muscle 4 in *w*^*1118*^ control and *Cln7*^*84D*^ larvae. A. The frequency (A) and amplitude (B) of miniature excitatory junction potentials (mEJP) are not significantly changed in *Cln7* larvae. mEJP frequency (left) was calculated over 120 s for each larva and amplitude (right) from 201-835 events using four different concentrations of extraceullular Ca^2+^. Mean is indicated by a solid bar; error bars indicate SD. 2-way ANVOA with Sidak’s multiple comparisons. F = 0.6086; DFn = 1; DFd = 66; p=0.4381 for both. B. Excitatory junction potentials (EJPs) are unchanged in Cln7 larvae. The mean of 16 EJPs at 1 Hz was calculated from n=13 (*w*^*1118*^) or 14 (*Cln7*^*84D*^) in 1 mM Ca^2+^. Mean is indicated by a solid line; error bars show SD. p=0.274; Mann-Whitney. C). *Cln7* mutant synapses display an increased rate of fatigue at 5 Hz stimulation. EJP amplitude at 5 Hz was normalised to the mean amplitude at 1 Hz stimulation immediately beforehand. n=4 NMJs for *w*^*1118*^ and =6 for *Cln7*^*84D*^. Trend lines fitted by linear regression are shown with solid lines with 95% confidence intervals indicated by dashes. R^2^: *w*^*1118*^ = 0.149; *Cln7*^*84D*^ = 0.383. The slopes are significantly different: F= 4.786; DFn = 1; DFd = 96; p=0.0311, t-test. D). Crawling speed is reduced in *Cln7*^*84D*^ larvae. Mean speed of *w*^*1118*^ control (n=109) and *Cln7*^*84D*^ (n=86) larvae was determined at 2.5 fps over 5 mins. **** p<0.0001; Mann-Whitney.

Nerve-evoked excitatory junction potentials (EJPs), were recorded at 1Hz frequency at 1 mM extracellular Ca^2+^ to evaluate synapse function. We found that EJPs at *Cln7* neuromuscular junctions were similar to those produced by control animals (Fig. 3B). We next applied continuous 5 Hz stimulation to the motor neurons to assess changes in the ability to recruit from the reserve pool of neurotransmitter vesicles. In the control animals, the evoked amplitudes followed a trend towards mild depression as vesicles in the reserve pool become depleted. However, a much more accentuated decline of EJP amplitude was observed in *Cln7* mutant animals under the same conditions (Fig. 3C). This suggests a potential defect in the ability of *Cln7* neurons to activate vesicles from the reserve pool or a defect in synaptic vesicle recycling.

The NMJs regulate body-wall muscle contraction. To identify if the electrophysiological changes we detected led to a quantifiable change in locomotion, we measured instantaneous speed of larval crawling. To address large variations in behavior within a population of larvae, and to avoid the tendency of larvae to avoid open fields, we used a custom 3D printed chamber to record the movement of multiple larvae simultaneously in oval “racetracks” over periods of 5 min (see Methods). *Cln7* mutant larvae move significantly more slowly than isogenic control larvae (Fig. 3D).

### Autophagy is unaffected in Cln7 mutants

Defects in the effectiveness of the autophagy pathway are associated with many forms of neurodegenerative disease, including forms of NCL (25) and impairment of autophagy is known to cause a reduction in synapse size similar to that seen in the *Cln7* mutants (21). In cultured cells, Cln7 is associated predominantly with lysosomes, which are essential mediators of autophagy and mutations affecting lysosomal function lead to a block in autophagic flux. However, in *Drosophila* normal lysosomal function seems to be critical in the neuron (19, 20, 26, 27), whereas Cln7 is restricted to the post-synaptic partner (13). Therefore, we asked whether impaired autophagy in the *Cln7* flies might underpin the synaptic phenotypes.

We examined accumulation of p62, a multiple domain protein that acts as a receptor to activate autophagy. Under nutrient-rich conditions, p62 is usually degraded in lysosomes but in starvation conditions autophagic flux is increased and some non-degraded p62 accumulates in lysosomes (Fig. 4A, see *w*^*1118*^ control). Thus, p62 levels are often monitored as one readout of autophagic flux. In *Cln7*^*84D*^ larvae no p62 accumulates under growth conditions. However, a small but not statistically significant amount of p62 did accumulate under starvation conditions in the *Cln7*^*84D*^ larvae, suggesting a minor impairment in autophagy or lysosomal function (Fig. 4A). This contrasted markedly with *atg18* mutants. Here, p62 clearly accumulated during growth and did so very dramatically under starvation conditions (Fig. 4A). Autophagy also functions normally in *Cln7* deficient cerebellar granule neurons (28).

**Figure 4.**
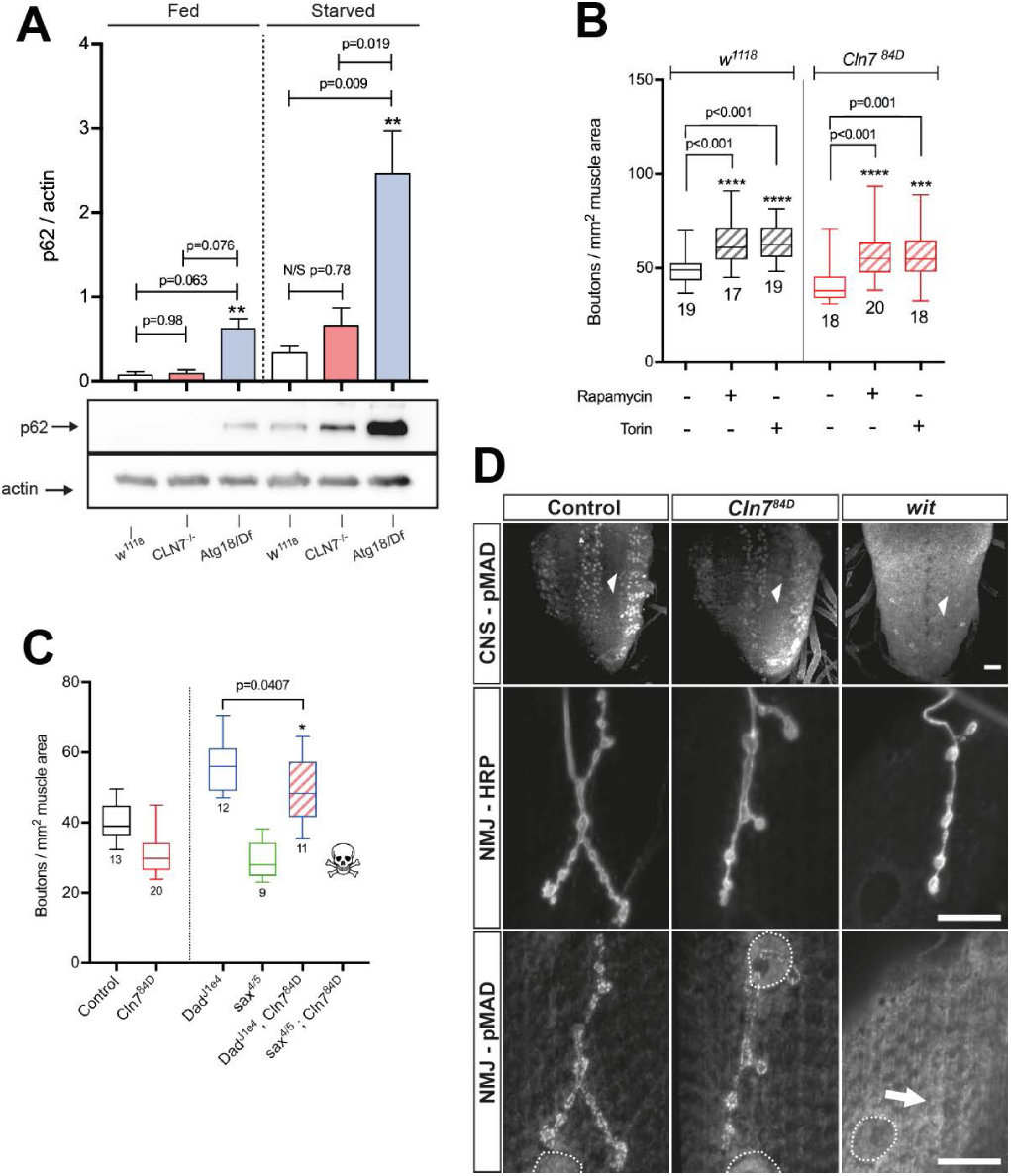
Autophagy and retrograde BMP signaling are unaffected by loss of Cln7. A-B). The lysosomal substrate, p62, does not accumulate in control of *Cln7*^*84D*^ larvae when feeding but does in *atg18* mutants. A. After 4 h starvation, p62 accumulates in control animals. Accumulation is slightly but not significantly increased in *Cln7*^*84D*^ larvae. In contrast, p62 accumulates to very high levels in starved *atg18* mutants. Bars show mean p62 band intensity +/- SEM normalised to actin, n=3 for each. ** p<0.01, ANOVA. C). Inhibition of TORC1 with Rapamycin or TORC1 and 2 by Torin drives NMJ overgrowth by increasing autophagic flux. *Cln7* mutant larvae respond similarly to control *w*^*1118*^ larvae suggesting no block in autophagy. *** p=0.01; **** p<0.001; ANOVA with Dunnett’s post-hoc comparisons, n is indicated. C). Mutations affecting retrograde signaling from the muscle alter neural development. Mutations in the inhibitory SMAD, *Dad*, drive overgrowth of the NMJ; mutations in the *saxophone* type I TGFβ receptor result in underdevelopment at a similar level to *Cln7* mutants. A double *Dad;Cln7* mutant displays a small but significant moderation of the *Dad* phenotype (2-tailed t-test: t=2.181, Df=21, p=0.0407). A *sax;Cln7* double mutant is lethal early in development. Control *w*^*1118*^ and *Cln7*^84D^ data from Figure are included for comparison. D). Mad is phosphorylated after binding of the Gbb ligand to Tkv, Sax and Wit receptors in the neuron. Characteristic pMad staining is clearly visible in columns of motor neurons in the control and *Cln7*^*84D*^ CNS but is absent in a *wit* mutant CNS (anti-pMad, top row, arrowheads). pMad staining in boutons is also present in boutons in control and *Cln7*^*84D*^ NMJs but absent in a *wit* mutant (anti-pMad, bottom column; anti-HRP marks the neuronal plasma membrane, middle column). Similarly, pMad is localised within muscle nuclei but absent in *wit* mutants (dashed circles). Scale bar = 20 μM.

We also examined whether increasing autophagic flux could rescue the *Cln7* phenotype by inhibiting the TOR complex activity using rapamycin or torin. TORC1 is a central regulator of growth and autophagy and homeostasis in cells and TORC2 is required for normal NMJ development (29). Inhibition of TORC1 increases autophagic flux which, in the *Drosophila* NMJ, leads to an overgrowth phenotype (21). Previously, it was reported that inhibiting TORC1 with rapamycin was unable to rescue the smaller NMJ phenotype seen in *atg18* mutants, presumably because autophagy is severely compromised in these animals and this cannot be circumvented (21). We reasoned that rapamycin treatment would similarly have no effect on the *Cln7* phenotype if it was due to defective autophagy. However, both rapamycin treatment or treatment with the TORC1 and TORC2 inhibitor, torin, caused a robust increase in NMJ size in both control and *Cln7* larvae indicating that autophagic flux can be increased (Fig. 4B). Taken together with the lack of p62 accumulation, and given that autophagy is not overtly affected in *Cln7* deficient cerebellar granule neurons (28), this suggests that the developmental changes in the *Cln7* animals are not due to defective autophagy.

### Normal BMP retrograde signaling in Cln7 mutants

Since Cln7 is required post-synaptically, we examined whether retrograde signaling necessary for normal growth of the pre-synaptic compartment of the NMJ might require Cln7 function. Retrograde signaling allows matching of pre-synapse size and excitability with muscle size; a key retrograde system is based on a BMP-type signal. Glass-bottomed boat (Gbb), a BMP molecule is secreted from the muscle and binds to Type I and II receptors Thick veins (Tkv), Saxophone (Sax) and Wishful thinking (Wit) expressed on the neuronal membrane. Ligand binding results in phosphorylation of the SMAD family member, Mothers against dpp (Mad) and its retrograde transport to the nucleus. Mutations in *gbb* or the receptors lead to loss of pMad and a smaller NMJ. Conversely, mutations in *Dad*, which encodes an inhibitory SMAD, lead to NMJ synapse overgrowth and a profusion of small, satellite boutons (19, 30-32).

Given enrichment of Cln7 at the PSD and the similarity of the *Cln7* mutant phenotype to those seen for mutations in components of the BMP signaling pathway, we asked whether BMP retrograde signaling is affected by loss of *Cln7*. Potentially Cln7 might act in the post-synaptic muscle cell in the production, processing or presentation of the Gbb ligand. In this case, loss of *Cln7* would give an identical phenotype to loss of Gbb, Tkv, Sax or Wit but a double mutant combination would not make the synapse undergrowth phenotype worse. Similarly, loss of *Cln7* would abolish overgrowth in *Dad* mutants by preventing Mad phosphorylation downstream of Tkv and therefore *Dad;Cln7* double mutants should have a *Cln7*-like undergrowth. We quantified NMJ size in *sax*^*4/5*^ and *Dad*^*J1e4*^ single mutants and saw the expected undergrowth and overgrowth phenotypes respectively (Fig. 4C). The NMJ synapses size in *sax* mutants was almost identical to the *Cln7*^*84D*^ mutants, however, a s*ax*^*4/5*^*;Cln7*^*84D*^ double mutant combination was lethal before late larval stage suggesting an additive effect and indicating that CLN7 and sax operate in different pathways. A *Dad*^*J1e4*^*;Cln7*^*84D*^double mutant combination showed a small reduction in overgrowth resulting in an intermediate phenotype. This suggests that phosphorylation of Mad in response to a Gbb signal is intact in the *Cln7* mutant larvae. To further confirm this, we stained the CNS of wild-type and *Cln7* larvae with anti-pMad. The nuclei of motor neurons clearly showed pMad present at the synapse in both genotypes (Fig. 4D).

### Loss of Cln7 leads to reduced TORC1 activity in the post-synaptic muscles

TOR complex activity is required for normal homeostatic mechanisms in the nervous system of both *Drosophila* and the mammalian CNS, including retrograde signals emanating from the post-synaptic cell (33-37). *Cln7* deficient MEFs show a defect in mTORC activation (9) so we considered whether dysregulation of TOR activity might underpin the reduced neural growth in the *Cln7* mutant flies. We stained NMJs with an antibody to phosphorylated S6, which is a marker of TORC1 activity (38). Phosphorylated S6 was seen clearly to localise at active zones in the pre-synaptic boutons (Fig. 5A). This pattern mirrors the localisation of phosphorylated p70 S6 kinase to active zones reported previously (39) and suggests localised activation of TORC1 at these release sites. The pS6 pattern was retained at active zones in the *Cln7* mutant synapses (Fig. 5G), indicating pre-synaptic TORC1 complex activity was normal, as expected.

**Figure 5.**
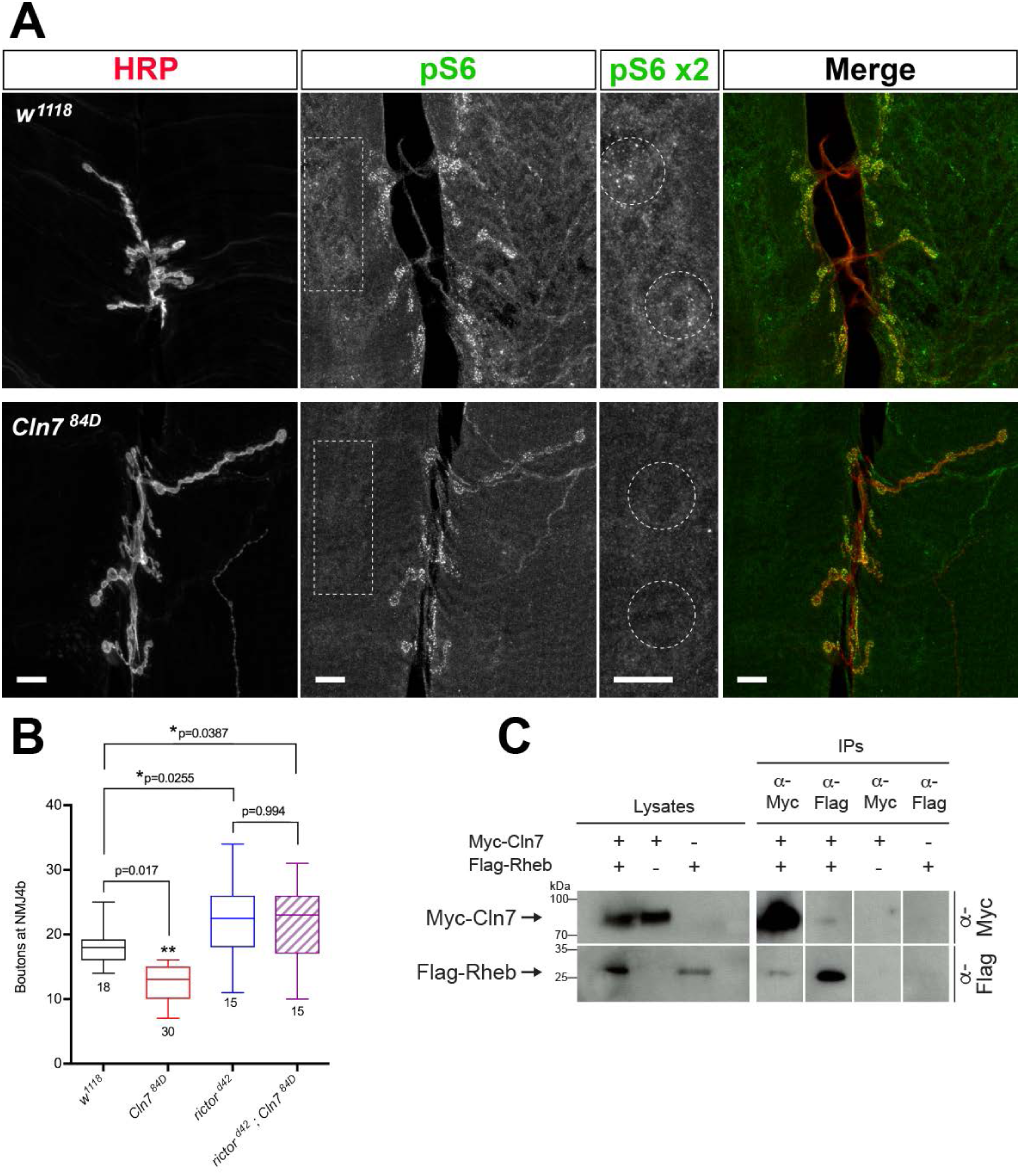
Loss of Cln7 leads to dysregulation of TORC. A). NMJ6/7 stained for the pre-synaptic membrane (anti-HRP, red) and a marker of TORC1 activity (anti-pS6, green). The RpS6 protein is phosphorylated by TORC1 activity via p70 S6 kinase. Distinct foci of pS6 are present at the active zones in pre-synaptic boutons in both *w*^*1118*^ *and Cln7*^*84D*^ larvae. pS6 staining is prominent in small vesicles in the post-synaptic muscle cell in *w1118* larvae close to the syncytial muscle nuclei. The pS6^+^ vesicles are largely absent in *Cln7*^*84D*^ larvae. Example regions of pS6 staining in the muscle are boxed and enlarged and perinuclear regions circled. Scale bar = 10 μm. B). TORC2 activity is required to restrict pre-synaptic growth. Flies mutant for the TORC2 component, *rictor*, have overgrown type 1b junctions. A *rictor;;Cln7* double mutant also shows an overgrowth phenotype. ANOVA with Dunn’s post-hoc comparisons, n is indicated. C). Cln7 can be immunoprecipitated in a complex with the TOR activating protein, Rheb. Myc-tagged Cln7 and Flag-tagged Rheb proteins were expressed in *Drosophila* S2 cells.

In the post-synaptic muscle cell strong pS6 staining was seen in vesicles that were primarily clustered near the syncitial muscle nuclei and likely to be lysosomes (Fig. 5A; dashed boxes are enlarged to the right, dashed circles in the enlargments show examples of clustered pS6^+^ vesicles). In contrast, the vesicular staining was reduced in the *Cln7*^*84D*^ mutant indicating reduced TORC1 activity in absence of Cln7 (Fig. 5A; dashed circles show equivalent peri-nuclear areas).

We next turned to TORC2. TORC2 acts in the neuron to restrict synaptic growth. A mutation in the TORC2-specific subunit, *rictor*, blocks TORC2 activity and leads to a large expansion of the NMJ (29). We first confirmed the *rictor* mutant phenotype and generated a *rictor*^*Δ42*^*;Cln7*^*84D*^ double mutant. In the double mutant, a similar expansion of the synapse occurred as in the *rictor*^*Δ42*^ single mutant (Fig. 5B). This places *Cln7* genetically upstream of TORC2 but is consistent with TORC2 function principally pre-synaptically in the neuron, downstream of Tsc2 (29), with Cln7 acting on the post-synaptic side.

We also identified biochemical evidence linking Cln7 with TORC1. Rheb is a small GTPase that activates mTORC1 in mammalian cells. We asked whether Cln7 might be present in the same protein complexes as Rheb. We expressed epitope-tagged forms of Rheb and Cln7 in *Drosophila* S2 cells and were able to co-immunoprecipiate the two proteins (Fig. 5C). Taken together, our data indicate that loss of *Cln7* leads to dysregulation of TORC1 activity in the post-synaptic cells. The consequences are developmental changes to the synapse and behavioural changes in the animal.

## Discussion

Here we have utilized *Drosophila* as a model system to identify an *in vivo* role for Cln7/MFSD8, the protein whose activity is reduced in late-infantile neuronal ceroid lipofuscinosis, an early onset childhood neurodegenerative disease. We demonstrate that Cln7 is required for normal synaptic development, consistent with a growing appreciation that the synapse is a significant target in neurodegenerative disease. When *Cln7* is disrupted, the *Drosophila* neuromuscular junction fails to reach its normal size having a reduced volume while maintaining normal active zone density. While this is characteristic of mutations affecting autophagic flux within neurons, autophagy is only mildly affected by loss of Cln7. Instead, Cln7 is resident in a vesicular compartment at the post-synaptic side of the junction where it functions to regulate TOR signaling. Previous studies have revealed that TOR signaling has a conserved postsynaptic role to influence retrograde regulation of synaptic activity at central and peripheral synapses in both *Drosophila* and vertebrates (36, 37) and that dysregulation of TORC signaling is associated with autism and cognitive decline (40). Recently it has also been observed that Cln7 is required for TOR reactivation in MEFs (9). We propose that late-infantile NCL may be a consequence of an early failure of synapse development brought about by dysfunctional TORC signaling.

### Lysosome protein function in neural development

Neuronal ceroid lipofuscinosis is a disease with multiple, overlapping forms and with varied ages of onset but which share lysosomal dysfunction as a shared feature. Lysosomal function is essential for autophagic flux and, consistent with a predominantly lysosomal localization for the Cln7 protein in cells, autophagy dysfunction is reported in a Cln7 mutant mouse model (10). Autophagy is also essential for neural development and long-term neural health, including in *Drosophila.* Given the similarity of the NMJ phenotype in *Cln7* and various *atg* mutants (21), we assumed defective autophagy was likely responsible for the *Cln7* phenotypes. However, autophagy appears to be required in the neuron rather than post-synaptically and, while we cannot rule out a small contribution, we demonstrate here that autophagy is largely intact in the absence of Cln7. Consistent with this, autophagy was also reported to be obstensibly normal in a *Cln7* deficient cerebellar granule neuron cell line (28).

Mutations in other lysosomal genes also affect NMJ development but these appear to act predominantly pre-synaptically in the neuron. Two notable examples are Spinster – like Cln7 an MFS-domain protein – which localizes to late endosomes and lysosomes in both neurons and muscle; and Trpml, a lysosomal cation channel and homolog of the gene mutated in the human lysosomal storage disorder, mucolipodosis type IV. In *spinster* mutants, a large overgrowth of the NMJ occurs due to excess reactive oxygen species being generated in the neurons (ROS are a cell-autonomous driver of neural growth in this system (26)). The retrograde BMP signaling that drives pre-synaptic expansion is also potentiated in the Spinster mutants, presumably due to inefficient degradation of the active ligand/receptor complex (19). In *Trpml* mutants, a similar undergrowth of the NMJ occurs to that in *Cln7* mutants (27). Trpml functions as a Ca^2+^ channel in lysosomes and by regulating an important aspect of lysosomal function – storage of intracellular Ca^2+^ – also regulates autophagy and TORC1 activity (20). However, like Spinster and in contrast to Cln7, Trmpl is required pre-synaptically. Here we demonstrate a post-synaptic specificity for Cln7 that exposes a potential new mechanism for regulating neural development and function in lysosomal storage disorders.

### TORC1 and TORC2 activity in neural development

Changes in synapse function have been seen in various models of CLN1 and CLN3 disease and may well be universal to the NCLs e.g. (15, 16, 41, 42). CNS pathology manifests very early in life in CLN7 disease so it is possible that developmental changes to the CNS and synaptic abnormalities (if also present in human CLN7 disease) predicate for later neuropathology, although potential mechanisms driving this are not yet clear. Might the failure to regulate TOR activity correctly underpin degeneration? The mTORC1 complex has a well-established role regulating neural growth and development in the mammalian nervous system. Several activating mutations upstream of mTORC1, including loss of PTEN or Tsc1/2, result in overgrowth of neurons and increase in density of dendritic spines, as do mutations in the translational repressor, FRMP, downstream of active mTORC1. Consistent with these effects, autism is a highly prevalent co-morbid feature in human disorders caused by mutations in these genes, including tuberous sclerosis, PTEN harmatoma syndrome and Fragile-X syndrome (40). TORC1 activity also has conserved functions regulating synaptic homeostasis (36, 37, 43). The effects of loss of TORC activity on metabolism – particular lipid metabolism – are well recognized (44) but how a reduction in TORC activity effects the nervous system, and how loss might contribute to neurological disease in the NCLs is less well understood. Our identification of a protein complex containing Cln7 and Rheb places Cln7 in the regulatory framework governing TOR activity in neurons but exactly how it functions will require identification of the other proteins in the complex and an understanding of the cargo transported by the Cln7 protein.

### Implications for the NCL

Lysosomal Ca^2+^ homeostasis is affected in several lysosomal disorders, including CLN3 disease (45) and, while Cln7 is not likely to be a Ca^2+^ channel, it will be interesting to ask if changes to Ca^2+^ accompany, and potentially underpin, the loss of post-synaptic TORC activity in Cln7 mutants, as they appear to pre-synaptically in Trmpl mutants (20). Many of the CLN proteins are considered to be lysosomal but lysosomes are predominantly localized in a peri-nuclear location in neurons. However, a proportion of Cln7 also localizes to the plasma membrane in non-polarised, cultured cells (6). This makes the presence of Cln7 in a vesicular compartment at the post-synaptic membrane intruiging, especially given the co-localisation of Cln7 and PSD-95 in the mouse retina (14). Palmitoyl protein thioesterase (Ppt1) is a lysosomal protease mutated in CLN1 disease. While Ppt1 is clearly lysosomal in most cultured non-neuronal cells, a synaptic pool appears to exist additionally in neurons (46). Potentially, alternative localizations for the CLN proteins *in vivo* in the CNS may be an important part of the emerging story of synaptic dysfunction in the NCLs.

## Materials and Methods

### *Drosophila* lines

*Drosophila* lines were obtained from the Bloomington Stock Center at Indiana University except for an isogenic *w*^*1118*^ control strain from the Vienna *Drosophila* Resource Center; PG-Gal4 (*NP6293) from the Kyoto DGRC Stock Center*; Spin-Gal4 and *Dad*^*j1e4*^ from Dr. Sean Sweeney (University of York, UK); and *rictor*^*Δ42*^ from Dr. Joseph Bateman (King’s College London). Gal4 lines used with stock numbers as of January 2019 were as follows: actin (BL3954); Elav^c155^ (BL458); Mef2 (BL27930); vGlut (BL26160); Repo (BL7415); PG (Kyoto: 105188). The Cln7 RNAi line used was yv;TRiP.HMC03819 (BL55664). To generate UAS-Venus-Cln7, the full-length Cln7 ORF was amplified by proofreading PCR using clone GH22722 as a template and cloned into pENTR (Thermo). After sequence verification it was recombined into the pTVW destination vector obtained from the *Drosophila* Genetics Resource Center, University of Indiana and transformants generated by BestGene.

### Generation of *Cln7* mutants

The P-element NP0345 inserted in the 5’UTR of the *Cln7* locus (CG8596) was excised by mating with to the Δ2-3 transposase line. Two independent alleles, 36H and 84D, were backcrossed over 6 generations to the isogenic *w*^*1118*^ control.

### Visualization of the larval NMJ

The pre-synaptic membrane was visulaised with Alexa-594 goat anti-HRP (1;1000; Jackson Immuno) except for volume measurements from confocal z-sections when a membrane-localized GFP was used (OK371-Gal4, UAS-CD8::GFP) with rabbit anti-GFP (AbCam, 1:1000). To ensure constant culture density and conditions between samples, cohorts of 100 1^st^ instar larvae were picked from the agar plates and transferred to vials of freshly made food and incubated for 3-4 days until wandering 3^rd^ instar stage. Larval flatmounts were immunostained using standard procedures, mounted in Prolong Gold (Thermo) and visualized on a Zeiss LSM510 Meta, LSM780 or LSM880 Airsycan confocal microscopes using using 40X N.A. 1.4 oil immersion, 40X N.A. 1.2 water immersion or 100x N.A. 1.46 oil immersion PlanApo lenses, respectively. Active zones were stained with mouse anti-Bruchpilot (clone NC82, 1:25; Developmental Studies Hybridoma Bank) and the post-synaptic density with anti-Dlg (clone 4F3, 1:20; DSHB). Rabbit anti-pMAD (1:500) was a gift of Prof. Carl-Henrik Heldin (Ludwig Institute for Cancer Research, Uppsala). YFP-Cln7 was visualized with rabbit anti-GFP (1:5000; AbCam ab290). Rabbit anti-pS6 (1:400) was a gift of Dr. Jongkyeong Chung, Seoul National University (38).

### Quantification of NMJ size

Volumes were acquired with an optical slice width of 1.0 μM and steps of 0.5 μM using a 40x N.A. 1.4 PlanApo objective and zoom = 2.0. Volocity was used to render the volumes then to measure the volume of the pre-synaptic membrane and to count active zones. Boutons were counted manually for the type Ib junction on muscle 4 of segments A3-A5. In each case a transmitted light image of the corresponding muscle was obtained at 10x and the muscle surface area calculated by outlining in ImageJ. Bouton number was then normalized to the muscle surface area.

### Quantification of larval movement

Wandering 3^rd^ instar stage larvae were transferred to a custom 3D-printed chamber comprising 12 oval troughs each 3 mm deep and with a track length of 75 mm. Each trough was lined with a thin coating of 3% w/v agar. The chamber was lit from below by a LED light table and filmed from above with a Basler GigE camera. Cohorts of 12 larvae crawling were filmed for 5 min each and their position tracked at 2.5 fps using Ethovision software (Noldus).

### Electrophysiology

Wandering 3^rd^ instar larvae were dissected in HL3.1 supplemented with Ca^2+^ concentration at 0.5, 0.75, 1.0 or 2.0 mM depending on the paradigm (47). Recordings were made from muscle 4 of segments A3-A5 using standard protocols (48). Baseline synaptic transmission was characterized by recording mEJPs for 2 min and 16 EJPs stimulated at 1 Hz. For high frequency stimulation, 16 EPSPs at 1 Hz were recorded followed by 5 Hz stimulation for 5 min. In all experiments a minimum of 8 larvae were used with maximum of 2 recordings per larva. In each case the starting resting potential of the muscle was −60 to −75 mV and resistance of 5-10 mΩ. Analysis of electrophysiological recordings was performed in Clampfit 10.0 (Molecular Devices) and data exported for further analysis in Prism 7.0 (Graphpad).

### *Drosophila* starvation assay

First instar larvae of control *w*^*1118*^, *Cln7*^*84D*^and *atg18a*^*KG03090*/^Df(3L)Exel6112 were transferred in cohorts of 20 from grape juice agar plates to standard food vials smeared with fresh yeast paste and allowed to develop for a further 24 h at 25 °C. Larvae were then removed from the food and protein samples prepared for each genotype. The remaining 10 larvae for each genotype were starved of amino acids for 4 h at 25 °C. Lysates of total protein were prepared by manually crushing the larvae with a mini-pestle directly into 200 μl Laemmli buffer containing 8 mM DTT and protease inhibitors.

### Cell transfection and immunoprecipitation

To generate expression plasmids, the *Cln7* pENTR clone was recombined into pAMW. The full-length Rheb ORF was amplified from clone GH10361 and cloned in to pENTR. After sequence verification it was recombined in pAFW. Tagged proteins were immunoprecipitated with Myc-Trap_MA beads (ChromoTek) or anti-FLAG M2 Affinity gel (Sigma) overnight at 4°C.

### Western blotting

Protein were separated by standard SDS-PAGE and transferred to PVDF membranes by wet transfer. Primary antibodies were: mouse anti-Flag M2 (1:5000; Sigma); goat anti-myc (1:1000; AbCam ab9132); rabbit anti-Ref(2)P (*Drosophila* homologue of p62, 1:500; a gift of Dr. Gabor Juhasz, Eötvös Loránd University, Budapest); rabbit anti-p70 S6K pThr389 (1:500; Cell Signaling Technologies) and mouse anti-actin (clone JLA-20, 1:500; DSHB). Secondary antibodies were HRP-conjugated goat antimouse or anti-rabbit IgG (both 1:4000, Cell Signaling Technologies). Detection of HRP was with SuperSignal West femto ECL reagent (Pierce) in conjunction with a Vilber Fusion FX system. p62 levels were normalized to actin from 16-bit images using ImageJ.

### Statistical analysis

All statistical tests were performed within Prism 7 (GraphPad). Normality of datasets was determined using a D’Agostino-Pearson test and datasets not conforming to Gaussian distributions were transformed with logarithmic or reciprocal transformations. Non-parametric tests were used where distributions remained non-Gaussian after transformation. Survival was compared by log-rank analysis. Samples sizes and tests used are described in the figure legends. Significance in all cases was set at p<0.05.

## Acknowledgements

The authors are very grateful to to Drs. Sean Sweeney and Joe Bateman for sending *Drosophila* lines; and to Dr. Gabor Juhasz, Prof. Carl-Henrik Heldin and Dr. Jongkyeong Chung for the anti-Ref(2)P, anti-pMad and anti-pS6 antibodies, respectively. Research was funded by The Wellcome Trust grant number 082004 to GT and by a UK Biotechnology and Biological Sciences Research Council New Investigator award BB/N008472/1 to RIT. BAS was supported by a Natural Sciences and Engineering Research Council of Canada grant number 250078. MO’H was supported by a King’s College, London Quota BBSRC PhD studentship and a KCL Peter Baker Travelling Fellowship. AM and KJC were supported by University of Birmingham College of Medical and Dental Sciences PhD studentship awards.

The authors declare no conflict of interest.

