## Supplementary figure 1 for "The neuronal ceroid lipofuscinosis protein Cln7 regulates neural development from the post-synaptic cell"

### Connolly et al
### Figure S1

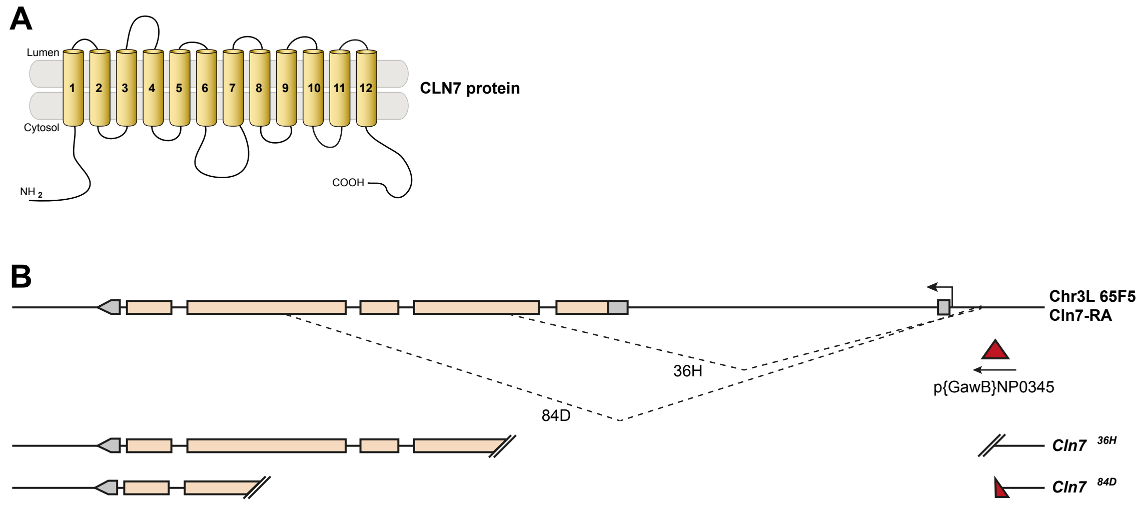


**Figure S1. Generation of *Cln7* mutations**

A. CLN7 is predicted to be a 12-span transmembrane protein and member of the multifacilitator superfamily of solute transporters.
B. P-element-mediated imprecise excision was used to generate deletions within the *Cln7* locus (CG8596). Two unidirectional deletion events, 36H and 84D, are shown diagrammatically. Some of the P-element sequence is retained in the 84D allele.
